# How context alters value: Price information recruits the brain’s valuation and affective regulation system for shaping experienced taste pleasantness

**DOI:** 10.1101/097915

**Authors:** Liane Schmidt, Vasilisa Skvortsova, Claus Kullen, Bernd Weber, Hilke Plassmann

## Abstract

Informational cues such as the price of a wine can trigger expectations about its taste quality and thereby modulate the sensory experience on a reported and neural level. Yet it is unclear how the brain translates such expectations into sensory pleasantness. We used multilevel mediation analysis of neural and behavioral data obtained in participants who tasted identical wines cued with different prices. We found that the brain's valuation system (BVS) in concert with the anterior prefrontal cortex explained the effect of price cues on taste pleasantness ratings. The sensitivity of the BVS to rewards outside the taste domain moderated the strength of these effects. Moreover, brain mediators of price cue effects overlapped with brain regions previously found to be involved in placebo analgesia. These findings provide novel evidence for the fundamental role that neural pathways linked to motivation and affective regulation play for the effect of informational cues on sensory experiences.

## INTRODUCTION

Past research has shown that information from the environment affects our value-related expectations and in turn how we report to evaluate experiences across a variety of sensory domains, including pain (Wager and Atlas, 2015), vision (Summerfield and de Lange, 2014), smell (de Araujo et al., 2005), hearing (Kirk et al., 2009) and taste (McClure et al., 2004; Nitschke et al., 2006; Plassmann et al., 2008). For example, informational cues such as the price of a wine induced expectations about its quality and how good it might taste; these expectations in turn modulated experienced taste pleasantness ratings (i.e., experienced value) and correlated with activation of brain regions involved in taste pleasantness encoding (i.e., the neural correlates of experienced value) (Plassmann et al., 2008).

However, it is not known whether the brain’s valuation system (i.e., the ventromedial prefrontal cortex and ventral striatum (Bartra et al., 2013; Clithero and Rangel, 2014; Levy and Glimcher, 2012) also *formally explains* changes in reported taste pleasantness in response to changes of value expectations. There are two streams of literature that support this hypothesis:

First, concepts in decision neuroscience show that the BVS encodes both expected and experienced value (Bartra et al., 2013; Platt and Plassmann, 2014; Rangel et al., 2008) and in turn might be crucial for translating price-based value expectations into experienced taste pleasantness.

Second, a meta-analysis of a related phenomenon in a different sensory domain—that is, how information about the efficacy of a treatment promotes pain analgesia (i.e., pain placebo effects) — showed that expectations induced by placebo cues led to decreased activity in pain-processing brain regions (Wager and Atlas, 2015). In addition, placebos also induced systematically across studies *increased* activity in the BVS. Although mediation has never been formally tested in these studies, these findings put forth the idea that value-related processes are an important antecedent causing expectancy effects of informational cues on sensory experiences to occur.

Against this background the goals of this paper are three-fold: First, we aimed at testing whether the BVS causally implements increases in experienced taste pleasantness ratings when higher prices generate higher value expectations. Importantly, a value-based modulation of experienced taste pleasantness ratings should be formally mediated by activity in the BVS, and if this is the case, these effects should be stronger the more sensitive an individual’s BVS is to reward.

Second, we also investigated whether the BVS is the only brain mediator underlying price expectancy effects or whether also other brain systems are needed to implement such effects. Prime candidate regions to support the BVS are brain regions in the lateral and anterior prefrontal cortex shown to be involved in the regulation of affective states (Ochsner et al., 2002, 2004) and stimulus values (Hutcherson et al. 2012)

Our third and last goal was to explore how generalizable our findings are across sensory domains by testing whether brain systems explaining price cue effects on taste pleasantness overlap with those explaining placebo analgesia. The idea behind this goal is based on findings from the neuroscience of placebo analgesia suggesting that descending pain modulation systems receive direct input from a set of brain regions including the BVS, the lateral and anterior prefrontal cortex and the amygdala (Wager and Atlas, 2015).

To investigate our hypotheses, participants took part in a previously used task assessing the effects of price cues on experienced taste pleasantness (Plassmann et al., 2008) (Figure 1a) while their brains were scanned using functional magnetic resonance imaging (fMRI). In this task, participants tasted three wines under different bottle price conditions (€3, €6, €18). Unbeknownst to the participants, real-world bottle prices were identical (€12). In some of the trials, participants received the wine sample for free; in other trials they had to pay a price proportional to the indicated bottle price. We were looking for neural systems that would satisfy three characteristics to link changes in price cues to changes in pleasantness ratings: they should (1) respond to price cues, (2) predict experienced pleasantness ratings and (3) explain the effect of price cues on experienced pleasantness ratings.

**Figure 1.**
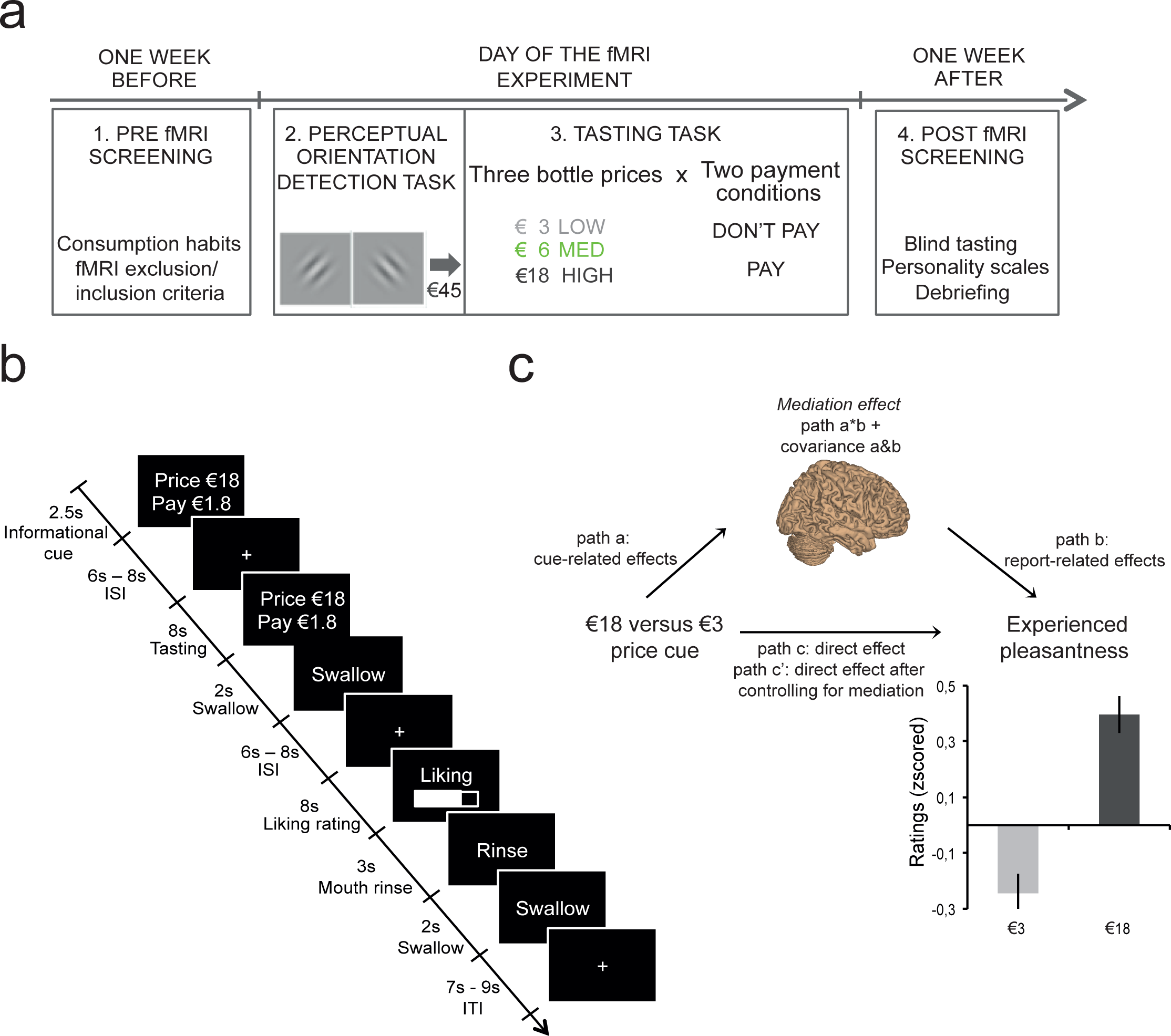
Experimental procedure, wine tasting task and results in n = 30 participants. **(A)** Experimental procedure. The experiment consisted of three events spread over two weeks. **(B)** Wine tasting task. Screenshots depict events within one trial with durations in seconds. Each trial started with the display of the price and payment condition. Following a jittered interstimulus interval (ISI) with a fixation cross on the screen, 1.25ml of wine was tasted via a tube and subsequently swallowed. Participants rated on a visual analogous scale their experienced pleasantness. They then rinsed their mouth with a water-like neutral liquid. Trials were separated by a jittered inter-trial interval (ITI) during which a fixation cross was displayed on the screen. **(C)** Mediation framework and behavioral results from N = 30 participants. Multilevel whole-brain mediation involved high and low price cue trials (€18, €3) as predictor variables, trial-by-trial beta images of brain activation at time of wine tasting as a mediator variable, and experienced taste pleasantness ratings as dependent outcome variable depicted in the bar graphs. Error bars correspond to the standard error of the mean (s.e.m.). Note that the effect of price cue was linear across all three price cue conditions, replicating prior work. There was no effect of payment condition, which was included into the model as a covariate of non-interest alongside wine type (see Figure S1 and Table S1).

To do so, we used the state-of-the-art analysis framework of multilevel brain mediation analysis (Wager et al. 2009; Wager et al. 2008). Mediation analysis advances inferences that can be drawn from standard univariate analyses in important ways. Standard univariate analysis allow either investigating the effect that an experimental condition such as a change in price cue has on neural activity or how neural activity correlates with reported change in sensory experiences. Yet, they don’t allow any inferences about the neural processes explaining the effect of price cue manipulations on pleasantness ratings. Mediation analysis goes beyond these two effects, by controlling for them and introducing a third variable - brain responses during wine tasting (Baron and Kenny, 1986). This approach thus creates a formal test of experimental manipulation-brain-behavior links (Atlas et al., 2010; Kenny, 2003; Lim et al., 2009; Wager et al., 2009).

We also leveraged the fact that some of our participants also took part in a previously used probabilistic monetary decision-making task (Fliessbach et al., 2010). In this task the participants could earn an additional payoff of 10 cents per trial by choosing one of a varying number of options on the screen. Importantly, the probabilities of receiving the additional payoff were known, so that no learning took place. The design of this task allowed us to capture how each participant’s BVS reacted to receiving a monetary reward. We used this measure as an estimate of individual differences in the sensitivity of each participant’s BVS outside the domain of taste rewards and thus independent from our main task. For these participants we then applied a multilevel whole-brain moderated mediation framework (Atlas et al., 2010; Woo et al., 2015) and tested whether individual differences of the BVS’s sensitivity in response to monetary rewards moderated price cue effects.

Lastly, we localized the brain mediators of price cue effects on experienced pleasantness within a set of brain regions reported to show increased activity in a meta-analysis of pain analgesia regions (Wager and Atlas, 2015).

## RESULTS

### Price cues increased experienced taste pleasantness ratings

First, we replicated the effect of price cues on experienced pleasantness ratings (Plassmann et al., 2008). As depicted in Figure 1c, we found that higher prices induced greater experienced pleasantness for identical wines (linear effect of price cue: β = 0.45, *SE* = 0.03, *p* < .001, 95% CI [0.39–0.51], Table S1). Our design also allowed us to explore whether such price expectancy effects depended on whether the participants had to pay for the wines they consumed. There was no significant main effect of payment condition (pay vs. consume for free) and no significant interaction between price and payment condition (Table S1, Figure S1). Moreover, we compared ratings influenced by price cues to baseline ratings that were sampled in a blind tasting a week after the fMRI session (M_blind_ = 5.03, SEM = 0.23). This comparison revealed that the price cue effects were driven by decreases in experienced pleasantness of low-priced (€3, M_informed_ = 4.19, SEM = 0.20) rather than increases in experienced pleasantness of high-priced (€18, M_informed_ = 5.21, SEM = 0.21) wines (paired, two-tailed *t*-test: €3 M_informed-blind_ = −0.83, SEM = 0.28 vs. €18 M_informed-blind_ = 0.18, SEM = 0.30, *t*(29) = −5.2, *p* < .001) (see SI paragraph 1.2.).

### Brain mediators of price cue effects on experienced pleasantness ratings

Second, we investigated which brain areas formally explained the effect of price cues on experienced pleasantness ratings. We conducted a multilevel whole-brain mediation analysis (http://wagerlab.colorado.edu/tools). As outlined in Figure 1c, the mediation analysis jointly considers two types of predictions at the time of wine tasting: (1) how price cues affect brain activity (path a) and (2) how brain activity predicts experienced pleasantness ratings (path b). Multilevel mediation is inferred if the direct effect from price to experienced pleasantness ratings (path c) gets significantly reduced after (path c’) controlling for the product of path a and path b coefficients within participants in addition to their covariance (cov (a,b)) across participants (Baron and Kenny, 1986). In the following we report the results for each of the three different paths (a, b and a*b+cov(a,b)) in the regression model:

#### 1) Price cue effect on brain activity at time of tasting (i.e., path a regression)

Path a assessed the price cue–related responses, which correspond to the relationship between price cue and brain activity. This effect is equivalent to the contrast high versus low price cue from standard univariate analyses (see SI paragraph 2.2.). As shown in Figure 2a, significant activations were found within the brain’s valuation system, including the ventromedial prefrontal cortex (vmPFC) and the bilateral ventral striatum (vStr), in line with previous findings (Plassmann et al., 2008). Other regions that displayed strong responses to tasting high-versus low-priced wines included the dorsolateral prefrontal cortex (dlPFC) and the lateral anterior prefrontal cortex (antPFC, BA 10) extending into the medial antPFC at a lower threshold; the primary gustatory cortex (i.e., insula); and semantic (Broca’s and Wernicke’s areas), motor, somatosensory and visual brain regions (precuneus and occipital cortex) (Table S2).

**Figure 2.**
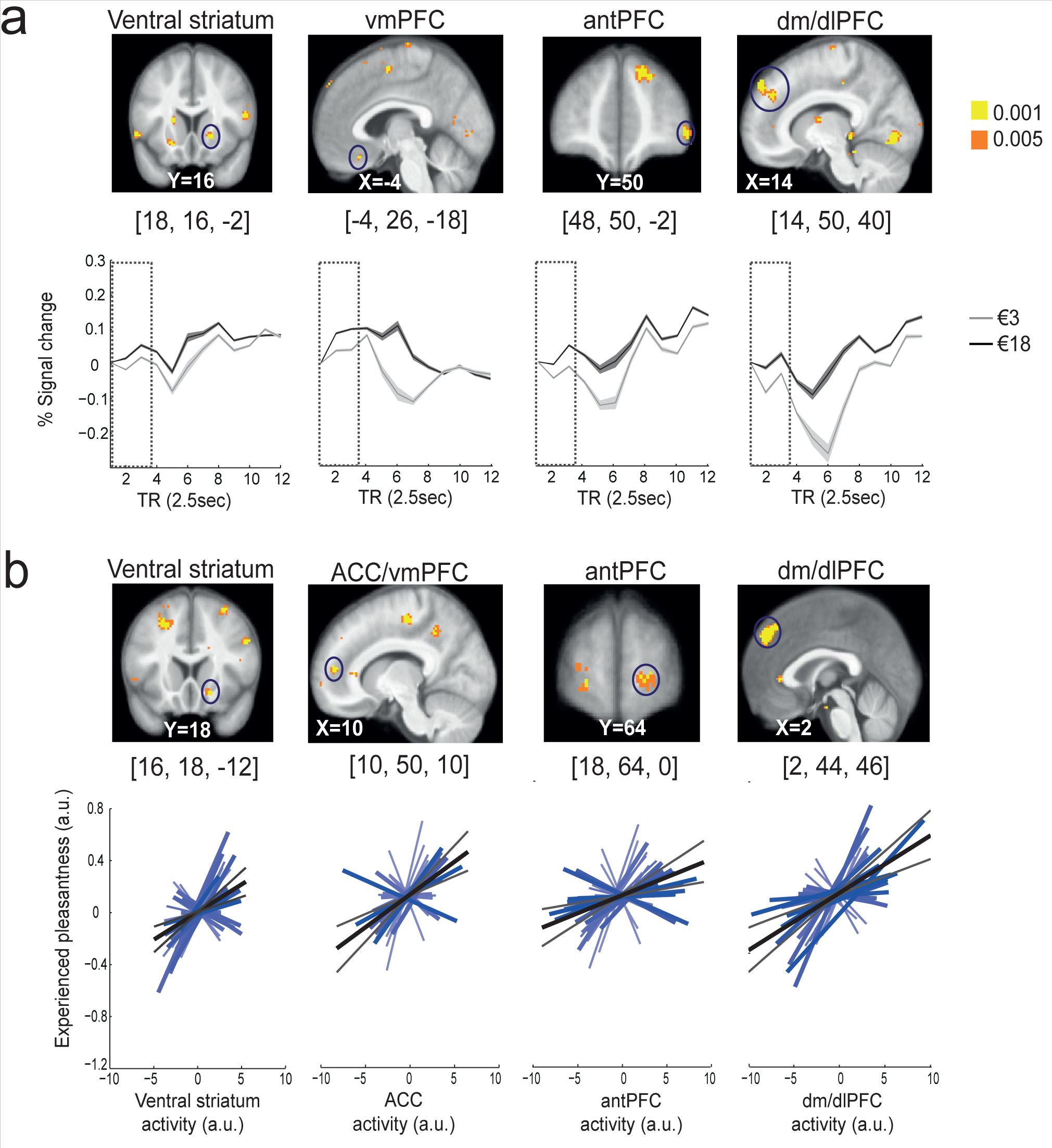
Brain responses to price cue and experienced pleasantness ratings averaged across n = 30 participants. Significant voxels are displayed in yellow *(p* < 0.001, uncorrected) and orange *(p* < 0.005, uncorrected) superimposed on the average anatomical brain image. The [*x, y, z*] coordinates correspond to Montreal Neurological Institute (MNI) coordinates and are taken at maxima of interest. **(A)** Path a: price cue–related effect. Activity in the ventral striatum, the ventromedial prefrontal cortex (vmPFC), the anterior prefrontal cortex (antPFC) and the dorsolateral/dorsomedial prefrontal cortex (dl/dmPFC) was greater following high price cues (€18) versus low price cues (€3). Line graphs depict time courses in each region of interest highlighted by blue circles. The dotted squares denote the wine tasting period. Shaded errors represent confidence intervals (means ± intersubject s.e.m.). Additional brain activations are listed in Table S2. **(B)** Path b: experienced pleasantness rating-related effect. Activity in the ventral striatum, the anterior cingulate cortex/ventromedial prefrontal cortex (ACC/vmPFC), the anterior prefrontal cortex (antPFC) and the dorsomedial prefrontal cortex (dmPFC) predicted parametric variation in experienced pleasantness ratings. Line graphs depict individual variations in path b coefficients (blue lines) with the group regression slope (gray line) within each region of interest highlighted by blue circles. Gray lines represent confidence intervals (95%). Additional brain activations are listed in Table S3.

#### 2) Brain activity predicting experienced taste pleasantness at time of wine tasting (i.e., path b regression)

Next, we looked for brain responses that predicted experienced taste pleasantness ratings. Importantly, this path b regression analysis controls for the effect of price cue on experienced taste pleasantness ratings. Thus, path b assessed brain activity that underpins endogenous variations in experienced taste pleasantness during wine tasting. We found that activity in the anterior cingulate cortex (ACC) adjoining the vmPFC, the right vStr extending into the nucleus accumbens (NAcc) and the hippocampus correlated significantly with variations in experienced taste pleasantness ratings irrespective of the price cue effects on these ratings (Figure 2b). Additional activation patterns were found in the lateral and medial part of the dorsal prefrontal cortex, and in more central regions in the bilateral anterior PFC (BA 10), somatosensory (posterior insula), middle temporal lobe, visual and motor cortex regions (Table S3).

#### 3) Brain mediators of price cue effects on experienced pleasantness (i.e., path a*b+cov(a,b) regression)

Third, we looked for brain regions that formally mediated the relationship between price cue and experienced taste pleasantness ratings. We found significant activations in the vmPFC, the right vStr and the anterior PFC (BA10) (Figure 3, Table S4) that satisfied the three criteria for formal mediation as outlined above(Baron and Kenny, 1986). Importantly, the regression coefficients for path a and path b were significantly correlated (*r* = 0.63, *p* < 0.01) for the vmPFC cluster, suggesting that its mediating role was driven by covariance. This is a common observation using multilevel whole-brain mediation analyses (Kenny, 2003). It implies that the vmPFC voxels consistently explained the effect of price cues on experienced pleasantness on the population level, although individual path a and path b coefficients varied in strength.

**Figure 3.**
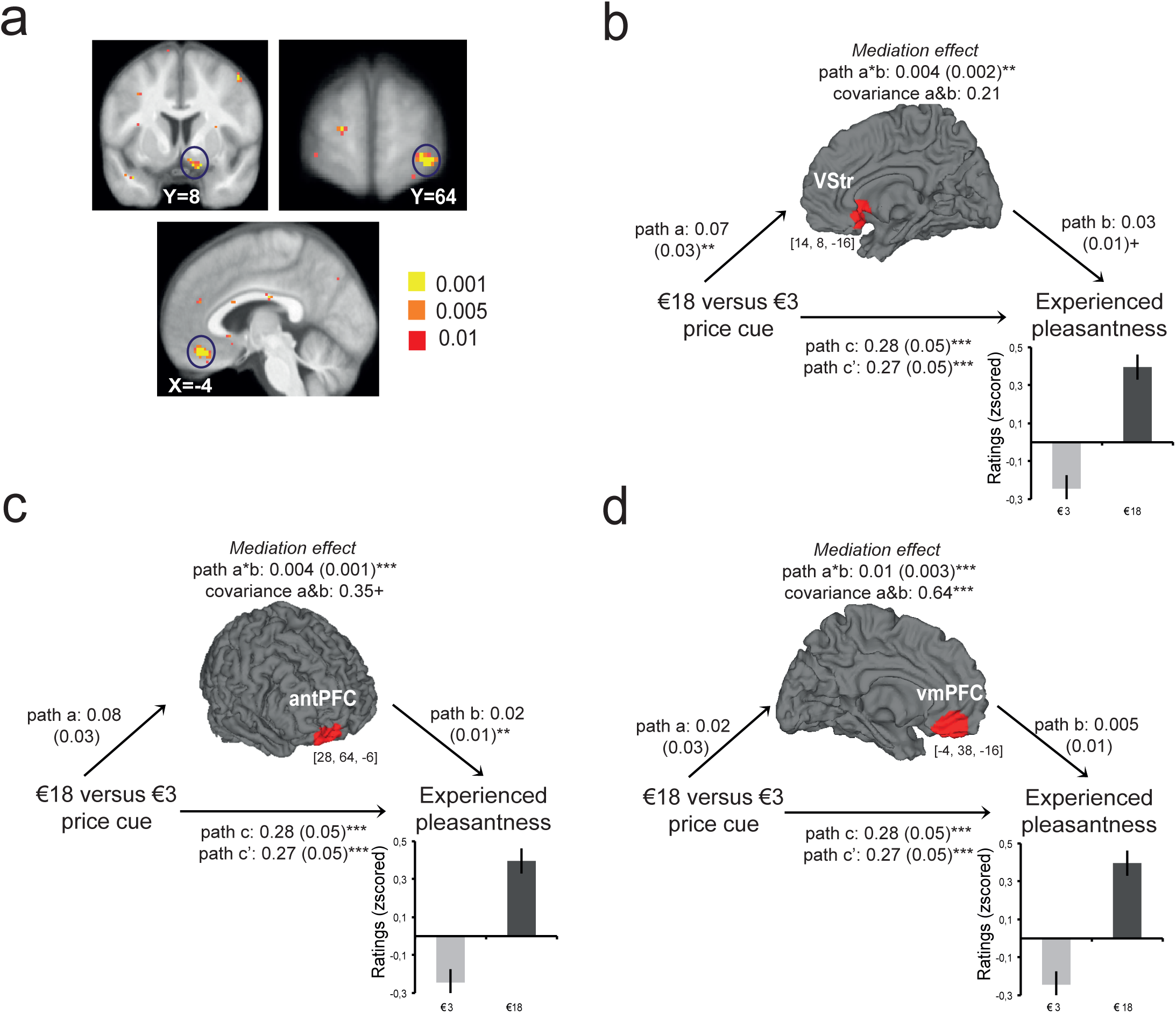
Brain mediators and moderators of price cue effects in n = 30 participants. **(A)** Statistical parametric maps of whole-brain activations mediating price cue effects on experienced taste pleasantness ratings. Significant voxels in yellow *(p* < 0.001, uncorrected) and orange *(p* < 0.005, uncorrected) are superimposed on the average anatomical brain image. Additional brain activations are listed in Table S4. **(B,C,D)** Multilevel mediation path diagram across N = 30 participants for the three brain mediators of price cue effects: (b) the ventral striatum, (c) the ventromedial prefrontal cortex (vmPFC) and (d) the anterior prefrontal cortex (antPFC). The [*x, y, z*] coordinates correspond to Montreal Neurological Institute (MNI) coordinates and are taken at maxima of interest. Average path coefficients (a*b (s.e.m.)) and the correlation of a&b coefficients (cov) across participants denote the joint activation in paths a and b at *** *p* < 0.001, ** *p* < 0.01 or ^+^ *p* < 0.05. Note that multilevel mediation effects can be driven either by significant path a and b coactivation or by covariance of path a and b coefficients. The three brain mediators of price cue effects were also located within a ROI mask of brain regions displaying increased activations under placebo analgesia (see Table S6).

Note that the path c’ regression assessing the direct effect of price cue on experienced pleasantness remained significant after controlling for activations of the brain mediators, suggesting a partial mediation role. Other brain mediators that were activated only by path a*b + cov(a,b) included temporal cortex, insula, and motor and visual brain regions (Table S4).

### General, task-independent sensitivity of the BVS moderates brain mediators of price cue effects on experienced pleasantness

To provide further evidence for the key role that the BVS plays in implementing price cue effects on experienced taste pleasantness, we investigated the role of individual differences in sensitivity of the BVS as assessed by its neural response to the receipt of monetary rewards in a different task. In line with our hypothesis, we found that neural activation of the BVS positively moderated price cue effects in the ventromedial prefrontal cortex and ventral striatum (Figure 4). Other path a–related brain activations moderated by individual sensitivity of the BVS involved the anterior prefrontal cortex, amygdala, dorsomedial prefrontal cortex, insula, periaqueductal gray and middle temporal gyrus (Table S5).

**Figure 4.**
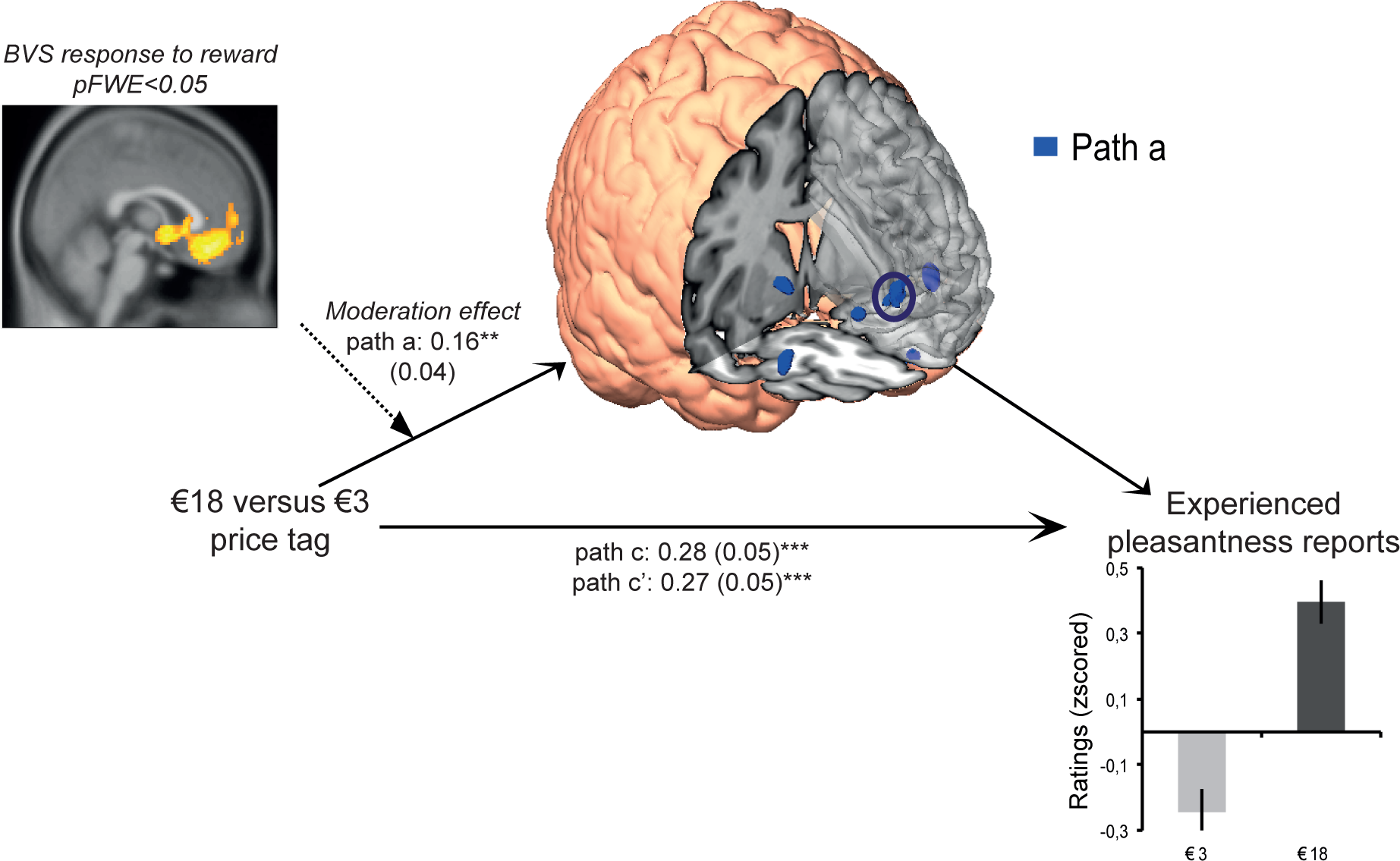
Whole-brain moderation of the price cue–related effect (path a) during wine tasting in a subset of n = 17 participants. The yellow and orange voxels depict significant brain activation by experienced reward during an independent monetary decision-making task, and are displayed at a threshold of *p*_FWE_ < 005 corrected for multiple comparisons on the cluster level (family-wise error). Blue voxels depict brain regions of the BVS that were positively moderated by the reward-related activation of the BVS during the monetary decision-making task. They are displayed at a threshold of *p* < 0.001 uncorrected at the voxel level. The path a coefficient corresponds (s.e.m.) to the moderated price cue effect on the vmPFC activation (^**^*p* < 0.01), highlighted by a dark blue circle. All voxels are superimposed on the average anatomical brain image. Additional path a related brain activation that was moderated by reward-related activation of the BVS during the monetary decision-making task is listed in Table S5.

### Localizing brain mediators or price cue effects on experienced taste pleasantness within brain regions of interest activated under placebo analgesia

Lastly, we localized brain mediators of price cues (i.e. path a*b+cov(a,b) activations) within a mask of brain regions of interest that were previously reported to display increased activation under placebo analgesia. We found that brain mediators in the anterior PFC, the nucleus accumbens/ventral striatum and the vmPFC were indeed located within this ROI mask (Table S6). This overlap of activations suggests the existence of a common neural signature for expectancy effects across sensory domains from pain to pleasure.

## DISCUSSION

The goal of this paper was to show that activity in the BVS plays a causal role in price cue effects on experienced pleasantness ratings. In this study we used a state-of-the-art methodological approach – that is multilevel, moderated mediation analysis providing novel evidence that (1) activity in the BVS formally mediated price cue effects and (2) individual differences in BVS sensitivity assessed in an independent monetary decision-making task moderated such price cue effects. Taken together, these two findings imply a robust and general key role that the BVS plays for such expectancy effects to occur.

An interesting question is whether both regions of the BVS (the vStr and the vmPFC) play the same role in price cue effects on experienced pleasantness. In several meta-analysis (Bartra et al., 2013; Clithero and Rangel, 2014; Levy and Glimcher, 2012) these two brain areas consistently and jointly correlated with the size of values assigned to different stimuli. The pain placebo literature (which is related to our question because it investigates the impact of informational cues from the environment on pain experiences) suggests, though, that they might play distinct roles: In their review Wager and Atlas proposed that the vmPFC might be an important hub to integrate all incoming information into “a coherent schema that informs and is informed by responses at other processing levels” (Wager and Atlas, 2015). This notion is in line with the idea that the vmPFC integrates different information into a valuation signal (Rangel et al., 2008). In addition, Wager and Atlas’s review also suggests that the vStr might be specifically linked to motivational processes during valuation—that is, a “wanting” to believe that one has received a painkiller does indeed translate into a less painful experience. This idea is in line with the linking of striatal activation to dopamine functioning that has been shown to be important for valuation (Schultz et al., 1997) and also for placebo effects (de la Fuente-Fernandez et al., 2001; Lidstone et al., 2010). Further research is needed to better understand the contributions of motivation and dopamine to expectancy-based placebo effects across sensory domains.

Our findings that the anterior PFC formally mediated price cue effects and that another brain region - the dlPFC - consistently activated in path a and b regressions provide first evidence for the recruitment of neural pathways involved in cognitive control of affective states for expectancy effects to occur. Functional magnetic resonance imaging studies from the field of social and affective neuroscience provided evidence that the anterior PFC and dlPFC are part of a set of frontal cortex regions that underpin the cognitive regulation of affective states (Ochsner et al., 2002, 2004; Levesque et al., 2003; Petrovic et al., 2005) and are associated with a variety of executive functions such as working memory (Gilbert and Burgess, 2008; Wager and Smith, 2003) and cognitive control in the sense of goal-directed action selection (Charron and Koechlin, 2010; Kouneiher et al., 2009). Notably, the anterior PFC is considered a “gateway” region that integrates incoming information from the environment with individual information from long-term memory (Gilbert and Burgess, 2008), supporting the idea that this system may implement expectancy effects of cues from the environment, such as the price of a taste sample or the white lab coat of the experimenter administering a placebo drug. One possibility could be that participants might recruit this brain region when reflecting on the external information from the price cue and their subjective beliefs and memories about how expensive and less expensive wines should taste. Additional evidence favoring this idea comes from a growing body of research on pain suggesting that frontal cortex brain regions involved in cognitive regulation processes overlap with brain regions associated with placebo analgesia (Benedetti et al., 2005). For example, the anterior PFC, and dlPFC are activated under the administration of a placebo drug in concert with verbal suggestion (Liebermann et al., 2004; Petrovic et al., 2005; Petrovic et al., 2002; Wager et al., 2004a; Wager et al., 2004b). Interestingly, the anterior PFC price cue mediator region indeed overlaps with a similarly located brain region activated under placebo effects of a sham anxiolytic drug (Petrovic et al., 2005). Yet the overlap of brain activation across different experimental contexts and tasks does not allow generalization of inferences about the specific underlying psychological processes (Poldrack and Yarkoni, 2015). We call for future studies that further test the causal contributions of cognitive processes such as emotion regulation and cognitive control for expectancy effects.

Finally, we explored whether the mediators of price cue effects on experienced taste pleasantness in the BVS and anterior PFC were located within a set of brain regions reported by a recent meta-analysis to activate under placebo analgesia (Atlas et al., 2010). We found evidence for an overlap between the same systems implementing expectancy effects in the pain and taste pleasantness domains. Thus, our findings provide first evidence for the idea that brain mechanisms implementing placebo effects might share common neural pathways across sensory domains. This first finding might help to reconcile previous debates contrasting whether there are variations in placebo responses (Benedetti et al., 2011) or whether they are similar (Wager and Atlas, 2015; Zubieta and Stohler, 2009). What our findings suggest is that there might indeed be differences in how the brain reacts to different contextual cues from the environment (i.e., the path a and path c responses) but that there might be shared neural pathways that translate those cues into experiences (i.e., that are mediating them). We call for future research that investigates this idea of shared neural pathways that causally implement expectancy effects across opposite sensory domains in more detail.

## AUTHOR CONTRIBUTIONS

H.P., V.S, & B.W. planned the experiment and developed the experimental design; C.K. collected the data; L.S. and V.S. analyzed the data under supervision of H.P.; L.S., H.P. & B.W. wrote the manuscript.

## ACKNOWLEDGEMENTS

We thank Ayelet Gneezy, Baba Shiv and Ab Litt for their help with the task design; Christina Walz for help with data collection; Tor D. Wager for scientific advice; and Lauren Atlas and Catherine Bushnell for sharing ROI masks. This study was funded by INSEAD’s research funds.

## SUPPLEMENTAL INFORMATION

The SI includes supplemental analysis and results with 6 figures and 11 tables. It can be found with this article online.

## EXPERIMENTAL PROCEDURE

### Participants

We recruited 54 healthy participants (21 male, 33 female; mean age = 29.1 years, SE = 1.1 years) via public advertisement at Bonn University. The study was approved by Bonn University’s Institutional Review Board. All participants gave written and informed consent before enrolling in the experiment. Participants were paid a show-up fee of €35 and received additional remuneration based on their performance in the study. Participants were screened for liking and at least occasionally drinking red wine. Standard fMRI inclusion criteria were applied to select participants. These included right-handedness, normal to corrected-to-normal vision, no history of substance abuse or any neurological or psychiatric disorder, and no medication or metallic devices that could interfere with performance of fMRI (SI Tables M7 and M8 for additional participant information). Twenty-four participants were excluded before the data was analyzed due to the following predefined exclusion criteria: head movement (≥3 mm; N = 17); incomplete data (N = 4) and insufficient orbitofrontal cortex coverage (N = 3), which was a priori defined as one region of interest based on previous findings and a meta-analysis on the brain’s valuation system (Plassmann, 2008, Bartra et al. 2013). Therefore, the analyses we used to investigate brain mediators of price cue effects on experienced taste pleasantness are based on a total of N = 30 participants (15 male, 15 female; mean age = 29.6 years, SE = 1.6 years).

In a follow-up analysis, we explored the role of individual differences in the sensitivity of the BVS, as sampled in an independent monetary decision-making task. To construct this individual difference measure (described in more detail below), we leveraged the fact that 17 of the 30 participants were also scanned with fMRI while performing a previously used monetary decision-making task (Fliessbach et al., 2010). Thus, the respective analyses were based on the subset of participants who took part in both tasks (i.e., 9 male, 8 female; mean age = 28.4 years, SE = 2.6 years).

### Procedure

The experiment consisted of three events spread over two weeks (Figure 1a). Participants were recruited via flyer advertisement for an fMRI experiment investigating how participants would evaluate wines under different conditions. The experimental procedure consisted of the following steps:

#### 1) Screening via phone interview (1 week before fMRI scanning).

During the phone interviews participants were screened for common fMRI exclusion criteria. Furthermore, we screened out wine experts and participants who did not like to taste wines or had dietary restrictions preventing them from doing so (see questions in Table M8).

#### 2) Perceptual orientation detection task (day of fMRI experiment before fMRI session).

Prior to scanning, participants performed a version of the Gabor orientation discrimination task previously used to test participants’ attention and perceptual learning skills (Stolte et al., 2014). The goal of this task was to implement a seemingly performance-based task that allowed us to have participants earn “house money” for the main tasting task, in which they were asked to spend some of the money they earned. In this Gabor orientation task participants were instructed to compare two Gabor patches and to decide whether they had the same or a different orientation without receiving feedback on the correctness of their responses. We used this task because it allowed us to adapt the task difficulty to each individual’s performance by varying the tilt. This was important because it allowed us to keep the performance and thus the performance-based payment constant for each participant. In other words, the task difficulty was adjusted such that the performance of each participant was around 60% to 65% correct answers. Every participant earned a total of €45 for his or her performance in the Gabor orientation task. The money was physically given to each participant after the task in a mix of coins and €5 bills in a bowl. The experimenter told the participants that the bowl would be kept in the fMRI control room and that money would be taken out each time the participant needed to spend money in the subsequent task.

#### 3) Wine tasting task

The wine tasting task was our main paradigm to investigate price cue effects on experienced taste pleasantness coding (Figure 1b). Participants repeatedly sampled 1.25 ml of three wines of the same retail price (€12). We had two experimental conditions: (1) We manipulated the bottle price information given to participants in each trial (i.e., €3, €6, €18), and (2) we manipulated whether participants could sample the wines for free or whether they had to pay for the sample a price that was 10% of the indicated bottle price (i.e., €0.3, €0.6, or €1.8). Importantly, each of the three wines was assigned to all experimental conditions over the course of the experiment, and we did not predict differences of our manipulations between the wines (i.e., wine was not treated as an experimental factor but was controlled for in all analyses reported below). Taken together, we applied a 3 (bottle price €3, €6, €18) x 2 (pay/no pay) within-participant experimental design. Each experimental condition was repeated 18 times (i.e., six times for each of the wines), resulting in a total of 108 trials. These trials were split up into three fMRI sessions of 36 trials with a duration of 30 minutes each. In total, participants consumed 135 ml of wine (less than a glass), and the total duration of the fMRI session was about 1.5 hours. Structural scans were acquired at the end of the fMRI session for 10 minutes. Screenshots of events within one trial are displayed in Figure 1b. Each trial started with the display of the price and payment condition information onset (2.5 s). A jittered inter-trial interval (ITI) (6 to 8 s) separated the price and payment information onset from the tasting period. During tasting the information about price and payment condition was again displayed on the screen, and the respective wine was delivered via an in-house–built electronic syringe pump system. The syringes filled with the respective liquids were placed on a MR-compatible system directly in the bore of the scanner to make the feeding tubes as short as possible. They were connected via a hydraulic tube system to electronic syringes in the control room. During this period, participants were instructed to swirl the liquid in their mouths (for a period of 8 s) and evaluate its pleasantness. They were also instructed to swallow only when the word “swallow” was displayed on the screen (2 s) to reduce head movement due to swallowing response as much as possible. After an ITI (6 to 8 s) participants were asked to enter their ratings of the pleasantness of the wine sample on a nine-point Likert scale from unpleasant to pleasant (8 s). After the rating, they rinsed their mouths with a neutral water-like liquid with a taste similar to saliva (containing 1g/l potassium chloride + 1g/l sodium bicarbonate + distilled water) (Plassmann et al., 2008) (3 s) and swallowed (2 s). Subsequent trials were separated by a jittered ITI (7 to 9 s) during which a fixation cross was displayed on the screen. Participants saw the information via goggles and indicated their responses using a response box system (both NNL, Bergen Norway). Bonn University’s in-house presentation software was used as experimental software to present the events and record responses.

#### 4) Blind tasting task (one week after the fMRI session)

One week after the fMRI session, participants performed a blind tasting evaluating pleasantness and taste of the three wines used during the fMRI session, without price or payment condition information present. They tasted 5 ml of each wine and rated pleasantness (one question: How much do you like this wine? not at all to a lot) and taste (two questions: How would you describe its taste? ordinary to extraordinary; inferior to superior) using a nine-point Likert scale. Participants were paid €10 for their participation in the blind tasting session.

### Monetary decision-making task

We used a monetary decision-making task that had effectively elicited activations in the brain’s valuation system in the past (Fliessbach et al., 2010). In this task, participants were instructed that their goal was to win as much money as possible by finding a circle in one out of a varying number of boxes displayed on the screen (see SI paragraph 2.3. and Figure M5). Importantly, this task was non-hypothetical and participants received additional payoffs based on their performance: That is, when participants guessed the right box as containing the circle, they were rewarded with a payoff of 10 cents; otherwise they won nothing. That means that each guess was associated with different known reward probabilities as a function of the number of boxes on the screen. In other words, the reward probability was 100% if only one box was shown, 50% for two boxes, 33% for three boxes, and 25% for four boxes. Because the reward probabilities were known, no learning was involved in this task. In total participants earned €5 for performing the task, and they made on average €4.61 (±€0.16) based on the number of correct guesses.

### Behavioral data analysis

All statistical tests were conducted with the Matlab Statistical Toolbox (Matlab 2015a, MathWorks). A linear mixed-effects model (using the *fitlme* function in Matlab) was fit for experienced pleasantness ratings with fixed effects for trial number (coded 1 to 36 per wine to correct for possible effects over number of tastings for each wine), wine (coded 1, 2, or 3 to test for differences in liking linked to the type of wine irrespective of price cue and payment condition), price (coded 1, 2, or 3), payment condition (coded 0 for free and 1 for pay), all possible two-way interactions, one three-way interaction trial by price by paying, and uncorrelated random effects for intercept grouped by subject (coded 1 to 30). All regressors were z-scored. Results are reported in Table S1. A similar analysis was performed for mean reaction times (SI Table M6).

### Image Acquisition

T2*-weighted echo planar images (EPI) with BOLD contrast were acquired on a 1.5T Siemens Magnetom Avanto scanner. We further applied a special sequence designed to optimize functional sensitivity in the orbitofrontal cortex. This consisted of a tilted acquisition in an oblique orientation at 30° to the AC-PC line. In addition, we used an eight-channel phased array coil that yields a significant signal increase in OFC over the standard head coil. To cover the whole brain with a repetition time of 2.5 seconds, we used the following sequence parameters: 31 axial slices; 3-mm slice thickness; 3 mm inter-slice gap. T1-weighted structural images were also acquired, co-registered with the mean EPI image, segmented and normalized to a standard T1 template, and averaged across all participants to allow group-level anatomical localization. The first three volumes of each session were discarded to allow for T1 equilibrium effects. Preprocessing consisted of spatial realignment, normalization using the same transformation as structural images, spatial smoothing using a Gaussian kernel with full width at half maximum of 8 mm, and high-pass temporal filtering (filter width 128 s). EPI images were analyzed using the Statistical Parametric Mapping software (SPM12; Welcome Department of Imaging Neuroscience).

### fMRI Analysis

We performed two analyses. In the first analysis, we were interested in identifying the formal brain mediators of price cue effects on experienced pleasantness ratings. In the second analysis, we were interested in whether the brain mediators from the first analysis were moderated by each participant’s BVS sensitivity, as measured by the neural response to monetary reward receipt during an independent monetary decision-making task.

#### 1) What were the brain mediators of price cues on experienced taste pleasantness ratings?

The analysis involved the following two steps:

##### a. Single-trial fMRI analysis

For whole-brain mediations we used a single-trial or “single-epoch” analysis approach (Buchel et al., 2002; Duann et al., 2002). Building on the implementation of the single-trial analysis for whole-brain multilevel mediations of cue effects on pain perception (Atlas et al., 2010; Woo et al., 2015) we applied this approach to the modeling of price effects on experienced pleasantness. A general linear model (GLM) design matrix estimated the magnitude of single-trial responses at the time of wine tasting (boxcar durations: 3 seconds) with separate regressors for each trial. Regressors of non-interest included “dummy” regressors coding for the intercept of each run, the linear drift across time within each run, six estimated head movement parameters from image realignment (x, y, z, roll, pitch, and yaw), their mean-centered squares, and their derivatives and squared derivative (total of 24 head movement regressors). Boxcar functions for each trial were convolved with the canonical hemodynamic response function. Note that estimations obtained by single-trial analyses are susceptible to noise from acquisition artifacts (sudden motion, scanner pulse artifacts, etc.). To assess the design-induced uncertainty due to collinearity with nuisance regressors we calculated variance inflation factors (VIFs). On average we excluded 1.06 trials (SD = 2.8) per participant (VIF ≥ 2.5). The VIFs of included trials ranged from 1.005 to 2.05. Single-trial beta images were then used as mediator variable *M* for the mediation analyses.

##### b. Multilevel whole-brain mediation analysis

We performed the multilevel mediation analysis using the Mediation Toolbox (http://wagerlab.colorado.edu/tools) (Wager et al. 2009; Wager et al. 2008) Mediation analysis extends standard univariate fMRI analyses by jointly considering the dynamic association between (1) experimental manipulation and brain activation and (2) brain activation and behavior. This is achieved by including a mediator variable *M* corresponding in the current study to brain activation at time of wine tasting. Under the null hypothesis of no mediation the two joint effects—price cue effect on brain activation and brain activation on experienced pleasantness ratings— are uncorrelated. Thus, a mediator variable is suggested to formally explain the covariance between a predictor variable *X* (i.e., price cue) on an outcome variable *Y* (i.e., experienced pleasantness rating). Taken together mediation analysis jointly tests three effects expressed by the following regression equations (Atlas et al., 2010):

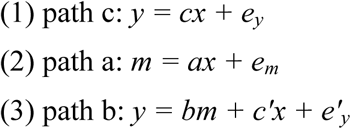

The variables *x, y*, and *m* correspond to trial by trial data vectors containing each subject’s experienced pleasantness ratings (*y*), price information (*x*), and data from each voxel at time of tasting (*m*). The variables e_y_, e_m_, and e’_y_ denote residual variance for each of the three regression analyses, respectively.

The first regression, called path c, corresponds to the direct (or total) effect of the experimental manipulation (*x* = price information) on behavior (*y* = experienced taste pleasantness ratings). The second regression, called path a, tests the relationship between the experimental manipulation (*x* = price information) on brain activity (*m* = activity in single-trial beta images). This effect is equivalent to the contrast high versus low price cue from standard univariate GLM analyses. The third regression, called path b, assesses the relationship between brain activity at time of tasting (m) and behavior *(y* = experienced pleasantness ratings) controlling for the experimental manipulation (*x* = price information). This effect, because it jointly controls for the effect of the experimental manipulation (i.e. price information), is also called report-related response identifying the brain regions that predict endogenously driven variations in experienced pleasantness. To control for additional experimental manipulations induced by wine type and payment conditions, these two variables were included into the mediation model as covariates of no interest.

Single-level mediation is assessed by the product of path a and path b coefficients: a * b = c – c’, with c’ corresponding to the direct effect of price tag on experienced pleasantness controlling for the brain activity at time of wine tasting (the mediator variable).

Importantly, the current study used a multilevel mediation analysis, which accounts for both within- and between-participant variations in one model by treating participant as a random effect. This involves testing on a first level the dynamic variations across trials within each participant between experimental manipulation, behavior, and brain activity, and on a second level for consistency of these variations across participants, allowing for population inferences. Crucially, multilevel mediation is inferred by the product of path a and path b coefficients and an additional component corresponding to the covariance of path a and path b estimates across participants (e.g., participants with strong path a effects also show strong path b effects) as expressed by equation 9 in Kenny et al. 2003:

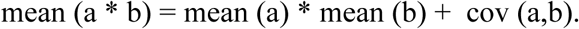

Thus, a multilevel mediation driven by covariance can identify voxels that consistently explain the effect of price cues on experienced taste pleasantness on the group level, although the path a and path b coefficients are heterogeneous on the individual level (e.g., negative for some and positive for others) (Atlas et al., 2010; Woo et al., 2015; Kenny et al.; 2003). We performed bootstrapping to test the significance of the path coefficients (Shrout et al., 2002; Efron et al. 1993). This involved estimating the distribution of individual path coefficients by randomly sampling with replacement 10,000 observations from the matrix of [a, b, c, c’, a*b] path coefficients. Two-tailed *p*-values were calculated from the bootstrap confidence intervals.

#### 2) Do brain mediators of price cues on experienced taste pleasantness ratings vary as a function of individual differences in the brain’s valuation system?

To test the moderating role of an individual’s neural sensitivity to experienced value, we extracted average beta estimates from 6mm-diameter spheres that were centered on the maxima of activation located in the ventromedial prefrontal cortex (vmPFC), the ventral striatum, and the adjacent anterior cingulate cortex (ACC), which activated significantly in response to the receipt of a monetary reward (i.e., experienced value) during the monetary decision-making task (see SI paragraph 2.3. and Table M5). The average of these beta estimates was used as a second-level moderator regressed to path a–related brain responses. We conducted the moderated, multilevel whole-brain mediation analysis in a subset of N = 17 participants.

#### 3) Do brain mediators of price cue effects on experienced taste pleasantness overlap with brain regions displaying increased activation under placebo analgesia?

To explore this idea we used a region of interest (ROI) mask that was previously reported by Wager and Atlas (2015). The ROI mask combined the vmPFC, dorsolateral prefrontal cortex, lateral orbitofrontal cortex, anterior prefrontal cortex, nucleus accumbens and ventral striatum, amygdala, hypothalamus, periaqueductal gray, and rostroventral medulla. It was used to mask mediating brain regions activating in path a*b+cov(a,b) at an uncorrected threshold of p<.001.

### Time courses

For all reported time course analyses, we extracted time courses from activations at maxima of interest. The response time courses were estimated using a flexible basis set of finite impulse responses, separated by one TR of 2.5 seconds.

## REFERENCES

Atlas, L.Y., Bolger, N., Lindquist, M.A., and Wager, T.D. (2010). Brain mediators of predictive cue effects on perceived pain. J Neurosci 30, 12964–12977.

Baron, R.M., and Kenny, D.A. (1986). The moderator-mediator variable distinction in social psychological research: conceptual, strategic, and statistical considerations. J Pers Soc Psychol 51, 1173–1182.

Bartra, O., McGuire, J.T., and Kable, J.W. (2013). The valuation system: a coordinate-based meta-analysis of BOLD fMRI experiments examining neural correlates of subjective value. Neuroimage 76, 412–427.

Benedetti, F., Mayberg, H.S., Wager, T.D., Stohler, C.S., and Zubieta, J.K. (2005). Neurobiological mechanisms of the placebo effect. J Neurosci 25, 10390–10402.

Benedetti, F., Carlino, E., Pollo, A. (2011). How placebos change the patient's brain. Neuropsychopharmacology 36(1), 339–354.

Buchel, C., Bornhovd, K., Quante, M., Glauche, V., Bromm, B., Weiller, C. (2002). Dissociable neural responses related to pain intensity, stimulus intensity, and stimulus awareness within the anterior cingulate cortex: A parametric single-trial laser functional magnetic resonance imaging study. J. Neurosci. 22, 970–976.

Charron, S., and Koechlin, E. (2010). Divided Representation of Concurrent Goals in the Human Frontal Lobes. Science 328, 360–363.

Clithero, J.A., and Rangel, A. (2014). Informatic parcellation of the network involved in the computation of subjective value. Soc Cogn Affect Neurosci 9, 1289–1302.

de Araujo, I.E., Rolls, E.T., Velazco, M.I., Margot, C., Cayeux, I. (2005). Cognitive modulation of olfactory processing. Neuron 46, 671–679.

de la Fuente-Fernandez, R., Ruth, T.J., Sossi, V., Schulzer, M., Calne, D.B., and Stoessl, A.J. (2001). Expectation and dopamine release: mechanism of the placebo effect in Parkinson’s disease. Science 293, 1164–1166.

Duann, J.R., Jung, T.P., Kuo, W.J., Yeh, T.C., Makeig, S., Hsieh, J.C., and Sejnowski, T.J. (2002). Single-trial variability in event-related BOLD signals. Neuroimage 15, 823–835.

Efron, B. and Tibshirani, R. J. (1993). An introduction to the bootstrap. Chapman and Hall/CRC.

Fliessbach, K., Rohe, T., Linder, N.S., Trautner, P., Elger, C.E., and Weber, B. (2010). Retest reliability of reward-related BOLD signals. Neuroimage 50, 1168–1176.

Gilbert, S.J., Burgess, P.W. (2008). Executive function. Current Biology 18, R110–114.

Hutcherson, C., Plassmann, H., Gross, J. Rangel A. (2012). Regulatory self-control involves a transfer of behavioral control from vmPFC to dlPFC valuation systems. Journal of Neuroscience, 32(39),13543–54.

Kenny, D.A., Korchmaros, J.D., Bolder, N. (2003). Lower level mediation in multilevel models. Psychological Methods 8, 115–128.

Kirk, K.I., French, B., Choi, S. (2009). Assessing spoken word recognition in children with cochlear implants. In Clinical management of children with cochlear implants, L.S. Eisenberg, ed. (San Diego, Plural Publishers), pp. 215–250.

Kouneiher, F., Charron, S., and Koechlin, E. (2009). Motivation and cognitive control in the human prefrontal cortex. Nat Neurosci 12, 939–945.

Levesque, J., Eugene, F., Joanette, Y., Paquette, V., Mensour, B., Beaudoin, G., Leroux, J.M., Bourgouin, P., Beauregard, M. (2003). Neural circuitry underlying voluntary suppression of sadness. Biological Psychiatry 53, 502–510.

Levy, D.J., and Glimcher, P.W. (2012). The root of all value: a neural common currency for choice. Curr Opin Neurobiol 22, 1027–1038.

Lidstone, S.C., Schulzer, M., Dinelle, K., Mak, E., Sossi, V., Ruth, T.J., de la Fuente-Fernandez, R., Phillips, A.G., and Stoessl, A.J. (2010). Effects of expectation on placebo-induced dopamine release in Parkinson disease. Arch Gen Psychiatry 67, 857–865.

Liebermann, M.D., Jarcho, J.M., Berman, S., Naliboff, B.D., Suyenobu, B.Y., Mandelkern, M., Mayer, E.A. (2004). The neural correlates of placebo effects: a disruption account. NeuroImage 2004, 447–455.

Lim, S.L., Padmala, S., and Pessoa, L. (2009). Segregating the significant from the mundane on a moment-to-moment basis via direct and indirect amygdala contributions. Proc Natl Acad Sci U S A 106, 16841–16846.

McClure, S.M., Li, J., Tomlin, D., Cypert, K.S., Montague, L.M., and Montague, P.R. (2004). Neural correlates of behavioral preference for culturally familiar drinks. Neuron 44, 379–387.

Nitschke, J.B., Dixon, G.E., Sarinopoulos, I., Short, S.J., Cohen, J.D., Smith, E.E., Kosslyn, S.M., Rose, R.M., Davidson, R.J. (2006). Altering expectancy dampens neural response to aversive taste in primary taste cortex. Nature Neuroscience 2006, 435–442.

Ochsner, K.N., Bunge, S.A., Gross, J.J., Gabrieli, J.D.E. (2002). Rethinking feelings: An fMRI study of the cognitive reguation of emotion. Journal of Cognitive Neuroscience 14, 1211–1229.

Ochsner, K.N., Ray, R.D., Cooper, J.C., Robertson, E.R., Chopra, S., Gabrieli, J.D.E., Gross, J.J. (2004). For better or for worse: neural systems supporting the cognitive down- and up-regulation of negative emotion. NeuroImage 23, 483–499.

Petrovic, P., Dietrich, T., Fransson, P., Andersson, J., Carlsson, K., and Ingvar, M. (2005). Placebo in Emotional Processing— Induced Expectations of Anxiety Relief Activate a Generalized Modulatory Network. Neuron 46, 957–969.

Petrovic, P., Kalso, E., Petersson, K.M., Ingvar, M. (2002). Placebo and opioid analgsia-imaging a shared neuronal network. Science 295, 1737–1740.

Plassmann, H., O’Doherty, J., Shiv, B., and Rangel, A. (2008). Marketing actions can modulate neural representations of experienced pleasantness. Proc Natl Acad Sci U S A 105, 1050–1054.

Platt, M., L. Plassman, H. (2014). Multistage valuation signals and common neural currencies. In Neuroeconomics, P. Glimcher, Fehr, E., ed. (Oxford, Academic Pressm Elsevier Inc.).

Poldrack, R.A., Yarkoni, T. (2015). From brain maps to cognitive ontologies: Informatics and the search for mental structure. Annual Review of Psychology 67, 587–612.

Rangel, A., Camerer, C., and Montague, P.R. (2008). A framework for studying the neurobiology of value-based decision making. Nat Rev Neurosci 9, 545–556.

Schultz, W. (1997). Dopamine neurons and their role in reward mechanisms. Curr Opin Neurobiol 7, 191–197.

Shrout, P. E. and Bolger, N. (2002). Mediation in experimental and nonexperimental studies: New procedures and recommendations. Psychol. Methods 7, 422–445.

Summerfield, C., and de Lange, F.P. (2014). Expectation in perceptual decision making: neural and computational mechanisms. Nat Rev Neurosci 15, 745–756.

Stolte, M., Bahrami, B., and Lavie, N. (2014). High perceptual load leads to both reduced gain and broader orientation tuning. Journal of Vision 14(9), 1–10.

Wager, T.D., and Atlas, L.Y. (2015). The neuroscience of placebo effects: connecting context, learning and health. Nat Rev Neurosci 16, 403–418.

Wager, T.D., Jonides, J., and Reading, S. (2004a). Neuroimaging studies of shifting attention: a meta-analysis. Neuroimage 22, 1679–1693.

Wager, T.D., Rilling, J.K., Smith, E.E., Sokolik, A., Casey, K.L., Davidson, R.J., Kosslyn, S.M., Rose, R.M., and Cohen, J.D. (2004b). Placebo-induced changes in FMRI in the anticipation and experience of pain. Science 303, 1162–1167.

Wager, T.D., and Smith, E.E. (2003). Neuroimaging studies of working memory: a meta-analysis. Cogn Affect Behav Neurosci 3, 255–274.

Wager, T.D., Waugh, C.E., Lindquist, M., Noll, D.C., Fredrickson, B.L., and Taylor, S.F. (2009). Brain mediators of cardiovascular responses to social threat: part I: Reciprocal dorsal and ventral sub-regions of the medial prefrontal cortex and heart-rate reactivity. Neuroimage 47, 821–835.

Wager, T. D., Davidson, M. L., Hughes, B. L., Lindquist, M. A., & Ochsner, K. N. (2008). Prefrontal-subcortical pathways mediating successful emotion regulation. Neuron 59, 1037–1050.

Woo, C.W., Roy, M., Buhle, J.T., and Wager, T.D. (2015). Distinct brain systems mediate the effects of nociceptive input and self-regulation on pain. PLoS Biol 13, e1002036.

Zubieta, J.K., and Stohler, C.S. (2009). Neurobiological mechanisms of placebo responses. Ann N Y Acad Sci 1156, 198–210.

